# Toward urban dark earths: Biochar-compost improves soil chemistry and reshapes dispersal and selection in community assembly

**DOI:** 10.64898/2026.11.26.769284

**Authors:** Nicholas Medina, Seneca Lee, Naim Edwards

## Abstract

Urban soils revitalized for agriculture invoke various management strategies from compost to potentially dark earth-inspired biochar. However, few studies analyze how biochar components holistically shape urban soil physical, chemical, and biological properties, including microbial community asssembly and potential functional groups. This study tested how compost co-application, biochar particle size, and application rate affect urban soil chemistry and ecological communities in Detroit, USA over two seasons. Results show that, 1) biochar alone lowered bulk density and increased bacterial dispersal limitation compared to compost alone, while biochar-compost together raised soil organic matter (SOM) and P and suppressed dispersal limitation; 2) particle size shaped microbial assembly, with fine particles <1 mm enriching topsoil copiotroph network modules, homogeneous selection in fungi, and potentially mites; and 3) higher application rates lowered bulk density, Ca, and cation exchange capacity, while raising SOM, P, and K, and favoring chemoheterotrophy and N cycling functions, shifting from oligotrophic to copiotrophic bacteria, but no change in richness. Overall, this study suggests that biochar-compost mixes improve urban soil chemical and biological fertility with some trade-offs.

## Introduction

Soils are degrading ten times faster than they form (Richter 2021), driven by industrial agriculture and intensive ur-banization (Beniston, Lal, and Mercer 2016), threatening regional food systems. Yet previously degraded urban soils re-purposed for agriculture after construction may regain fertility relatively quickly (Mukherjee, Lal, and Zimmerman 2014) while far from carbon (C) saturation (Six *et al*. 2002; Stewart *et al*. 2007, 2009), with recovery depending on specific function and multi-functionality responses to diverse management strategies (Stratton *et al*. 2022; Edwards, Medina, and Asker 2023). Beyond traditional fertilizer and compost additions to vacant land, industrial pyrolysis of biomass for soil amendment, or biochar, has emerged as a technology modeled on global Indigenous dark earth soils (*terra preta de indio*) made using sustainable waste management strategies (Woods *et al*. 2009; Silva *et al*. 2021, 2022; Asare 2022; Lombardo *et al*. 2022) that outperform surrounding soils (Glaser 2007). Biochar has the potential to offset 12% of industrial C emissions (Woolf *et al*. 2010), and while its effects remain context-dependent (Liu *et al*. 2016; Joseph *et al*. 2021), meta-analyses show higher benefits in soils low in organic C and cation exchange capacity (Crane-Droesch *et al*. 2013; Jeffery *et al*. 2017), yielding up to 14 times more C and nitrogen (N) in nutrient-poor compared to nutrient-rich soils (Deng *et al*. 2026). As a result, biochar amendments may be especially useful for young vacant lots (Midgley, Anderson, and Minor 2021).

Biochar effects on young urban soils may differ from those of compost alone, including by application rate and material particle size. Compost improves various associated benefits, including added phosphorus (P) and other nutrients (Small *et al*. 2019) and lowering bulk density (Singh *et al*. 2022). In comparison, biochar may yield similar benefits, but by retaining P and Ca rather than adding nutrients (Lombardo *et al*. 2022), as well as higher surface area and irregular particle shapes (Quilliam *et al*. 2013; Jaafar, Clode, and Abbott 2015) mediating structural benefits (Wang *et al*. 2017), such as water retention (Liu *et al*. 2017; Liao & Thomas 2019; Edeh & Mašek 2022; De Pauw *et al*. 2023). In practice, higher application amounts (10-20% vs. 1-5%) may yield faster (Verheijen *et al*. 2019) yet saturating (Liu *et al*. 2016) and more costly benefits (*ca*. 3-5 USD m^-2^ [0.3-0.5 USD ft^-2^]) compared to lower application rates (1%) with delayed benefits.

As soil amendment strategies shape microbial hotspots (Kravchenko *et al*. 2019) they then likely affect soil microbial communities (Anderson *et al*. 2014; Chen *et al*. 2017; Lévesque *et al*. 2020; Han *et al*. 2022), such as P-scavenging arbuscular mycorrhizal fungi (Warnock *et al*. 2007), as well as the underlying processes behind how their biodiversity assembles (Vellend 2010; Nemergut *et al*. 2013; Ning *et al*. 2020). Compost, higher activated biochar application rates, and smaller biochar particle sizes may enrich copiotrophic species favored under high nutrients such as N-cycling groups (Anderson *et al*. 2014), and/or facultatively anaerobic groups in low-oxygen pockets under very high application rates. This may occur due to homogeneous selection for specific metabolic capacities, or heterogeneous selection for any anoxic energy acquisition strategies. Such shifts may improve or lower overall soil health, as enriched copiotrophs and N-cyclers may increase nutrient availability, but also accelerate soil organic matter (SOM) turnover and nutrient losses from leaching (Shiu *et al*. 2022; Feng *et al*. 2023). Higher trophic levels in soil food webs may also respond, such as earthworm avoidance of small biochar particle sizes (Prodana *et al*. 2019), but invertebrate grazer responses appear inconsistent and understanding remains limited (Domene 2016). Overall, biochar effects on soil biota remain context-dependent (O’Neill *et al*. 2009; Noyce *et al*. 2015; Lucheta *et al*. 2017; Brtnický *et al*. 2021), and few studies analyze responses of re-purposed urban soils under land-use turnover (Bray & Wickings 2019; Nugent & Allison 2022; Schröder *et al*. 2024).

This study tested how compost co-amendment, biochar particle size, and application rate affect SOM and nutrient availability, as well as microbial and invertebrate diversity, assembly, and potential ecological functions in a recently re-purposed urban agricultural site. It was hypothesized that 1) biochar-compost would differ from either amend-ment alone, because compost supplies nutrients and microbial biomass while biochar supplies pore habitat, shifting microbial and invertebrate composition, assembly, and functional potential; 2) small biochar particles would change soil properties and biota more than large ones, by adding more surface area and filling more pores; and 3) more biochar would most lower bulk density and raise SOM and nutrients, via more niche space for microbial diversity.

## Methods

### Study site

This study was conducted on a 225 m^2^ (2500 ft^2^) research parcel at the Michigan State University Extension Detroit Partnership for Food Learning and Innovation (MSU-DPFLI, Detroit, MI, USA; 42.4124° N, 83.2630° W). The area was minimally amended since a school was demolished in 2016, and mapped as Brems-Urban land complex (mixed, mesic Aquic Udipsamments; loamy sand fill over glaciolacustrine sands; Soil Survey Staff 2024), with topsoil A horizons <5 cm deep, and a humid continental climate (10.1 °C and 864 mm yr^-1^; Hart & Bell 2015). Baseline soils had heavy metals below legal concern thresholds, 2-3% SOM, pH 8, and aggregate stability <22%. A previous study on this site showed that roto-tilling reduced penetration resistance and improved infiltration, with tailored mixed cover crops able to suppress weeds (Edwards, Medina, and Asker 2023).

### Study design

The study area was initially roto-tilled to 10 cm depth, subsequently only weed-trimmed, and divided into 64 adjacent 2.25 m^2 plots separated by 30 cm wide wood-chip and paper-mulch paths and grouped into blocks with at least four replicates per treatment. Biochar-containing treatments used pine wood biochar (*ca*. 0.76 m^3^ [1 yd^3^]; Wakefield Biochar, Columbia, MO, USA; southern yellow pine pyrolyzed at >700 °C, 85-95% C, ∼5% ash, pH 8-9, 0.1-2 mm particles, ∼365 m^2^ g^-1^ surface area), which was hand-sieved at 1 mm over two weeks into small (<1 mm) and large (>1 mm) particles, mixed 1:1 by volume (v/v) with local compost (Tuthill Farms & Composting Inc, South Lyon, MI, USA), and was activated outside 48h before application.

In the first season, all 64 plots (16 blocks of four) received compost alone (‘compost’) or unsieved (‘unsieved’), small (<1 mm), or large (>1 mm) biochar-compost (*n*=16 plots each) at 5% v/v of the top 10 cm (150 x 150 x 10 cm) two days before seeding (hereafter the ‘particle size’ treatment category). Plots were hand-sown with a hoe at recommended spacing in two alternating rows per species of cowpea (*Vigna unguiculata*), a legume complementing nitrogen-poor biochar and culturally important to local farmers, and fast-growing, P-scavenging, pollinator-attracting buckwheat (*Fagopyrum esculentum*; eight buckwheat and five cowpea seeds per row, *ca*. 16 and 10 per plot; Keep Growing Detroit, MI, USA; Johnny’s Selected Seeds, Albion, ME, USA; Clark 2007). In the second season, one quarter of plots kept particle size treatments (*n*=4 each); one quarter received unamended (‘none’), biochar-only (‘biochar’), compost-only (‘compost’), or mixed-size biochar-compost (‘both’) treatments (*n*=4 each) to separate direct biochar effects from compost co-application (‘*vs*. compost’ treatment category); and half received biochar-compost with 0, 5, 10, or 20% biochar v/v (*n*=8 each) as a *ca*. 1-2 cm surface layer (Ryals *et al*. 2016) to test maximal soil responses (‘amount’ or application rate treatment category). The second spring, a red clover (*Trifolium pratense*) and hairy vetch (*Vicia villosa*) cover crop mix was broadcast, followed by beet (*Beta vulgaris*) and field pea (*Pisum sativum*) in four alternating rows of eight to ten seeds each 2.5 cm deep, which did not germinate.

### Soil sampling

Soil cores (5.5 × 10 cm metal corer with a sand-filled dead-blow hammer for minimal disturbance; AMS Inc, American Falls, ID, USA) were collected in triplicate per plot in autumn of year one and in duplicate in spring after year two, split into 0-5 and 5-10 cm layers along a natural horizontal fault near 5 cm, and stored at 4 °C within 24-72 h (320 soil samples). Cores were not pooled: all were used for bulk density, the first core per plot for chemistry, and the second for DNA (128 samples each). Soil pH (1:1 soil-water), Olsen P (Olsen *et al*. 1954), and ammonium-acetate extractable potassium (K), magnesium (Mg), and calcium (Ca) were measured at the Michigan State University Soil and Plant Nutrient Laboratory (East Lansing, MI, USA), which also reported SOM via loss on ignition and cation exchange capacity (CEC; sum of extractable K, Mg, and Ca). Nine Bray P1 samples were excluded.

Soil DNA was extracted from 0.15 g soil with PowerLyzer PowerSoil kits (QIAGEN), modified with a 1:1 mix of PowerBead solution and 1 M sodium-potassium phosphate buffer (pH 7, 300 µL), a 10-min 65 °C incubation, and a 5-min -80 °C freeze to improve desorption from fine particles. Bacterial 16S rRNA V4 (515F/806R; Caporaso *et al*. 2011) and fungal ITS2 (5.8S-Fun/ITS4-Fun; Taylor *et al*. 2016) amplicons were subject to paired-end sequencing (Kozich *et al*. 2013) on an Illumina MiSeq at the University of Michigan Microbiome Sequencing Core (Ann Arbor, MI, USA). Ground-dwelling invertebrates were collected in year one with a soap-water pitfall trap per plot over one week, stored in 70% ethanol, and identified to order and morphospecies.

### Bioinformatics

Higher quality forward reads were processed using *DADA2* v1.28 (Callahan *et al*. 2016) (truncated to 240 and 160 bp; max. expected errors 2 and 5; merged) and ITS forward reads (*cutadapt* ; Martin 2011; Rolling *et al*. 2022; max. expected errors 8; ≥50 bp) after consensus chimera removal, classified (Wang *et al*. 2007) against SILVA v138.1 (Quast *et al*. 2012; Glöckner *et al*. 2017; McLaren & Callahan 2021) and UNITE v9.0 (Abarenkov *et al*. 2010; Nilsson *et al*. 2019; Abarenkov *et al*. 2022). Samples, with median 2,402 bacterial and 2,592 fungal reads, were rarefied to 1000 reads to maintain treatment representation (Schloss 2024; Good’s coverage 95.8±0.3% and 98.9±0.1%). Bacterial functions (FAPROTAX; Louca, Parfrey & Doebeli 2016) and fungal lifestyles (FungalTraits; Põlme *et al*. 2020) were assigned at the finest resolution (Toole *et al*. 2021), merging nested groups (aerobic into chemoheterotrophy, nitrogen respiration into nitrate reduction), with *microeco* (Liu *et al*. 2021) and *meconetcomp* (Liu *et al*. 2023).

### Statistics

Data were analyzed separately per treatment category in R 4.6 (R Core Team 2026). To facilitate comparisons across response variable scales, several metrics were normalized to treatment means as z-scores: including bulk density, chemistry and nutrients, richness, the six most abundant bacterial phyla and eight most abundant fungal orders, functional groups ≥0.5%, and abundant invertebrate orders, which were analyzed using one generalized additive model across treatments (GAM; *mgcv* ; Wood 2011) with *gratia::appraise()* residuals checks (Simpson 2024) and confirmed with analogous linear mixed model formulations (*glmmTMB*; Brooks *et al*. 2017). Models tested treatment × depth, smoothed application rate by variable with curvature *k* =3 with random block and plot intercepts, plus year for particle size treatments. Post hoc contrasts (*emmeans*; Lenth 2026) were reported for significant treatment effects with Benjamini-Hochberg (BH) corrections (*multcomp*; Hothorn, Bretz & Westfall 2008). Microbial and invertebrate composition was tested with PERMANOVA (*vegan*; Dixon 2003; *R*^2^; 9999 permutations of samples permuted among plots within blocks, and years for particle size) and dispersion similarly with PERMDISP (Anderson 2006), ordinated using distance-based redundancy analysis (db-RDA) with *envfit* soil and functional-group vectors.

Bacterial and fungal assembly processes were partitioned with *iCAMP* (Ning *et al*. 2020) using phylogenetic bins of a neighbor-joining tree (*phangorn*; Schliep 2011) of *MAFFT* -aligned ASVs (Katoh & Standley 2013), relatedness, and Raup-Crick null models, and resulting process importance metrics were modeled similarly as above. A taxonomy-bounded hybrid fungal tree was used to improve ITS alignment with *DECIPHER* (Wright 2016). Genus abundances were used to construct a signed co-occurrence network (*igraph*; Csárdi & Nepusz 2006) using genera in ≥20% of samples linked by within-year Spearman correlations of centered log-ratio (CLR; Aitchison 1986) transformed counts, keeping both positive and negative correlations with absolute value strength ≥0.3 and false discovery rate-adjusted *p*<0.05. Leiden modules (Traag, Waltman & van Eck 2019) were identified using positive edges, where modularity (*Q*) and clustering (*C*) were compared with 999 random networks, the proportion of negative edges between modules compared to 999 module-label permutations, and degree distributions were tested for power laws (*microeco, poweRlaw* ; Clauset, Shalizi & Newman 2009; Gillespie 2015). Module abundances and sample sub-network properties (*meconetcomp*), including robustness to random node removal, were modeled as above. Differential abundance of the most abundant genera and responsive network modules was tested with *DESeq2* (Love, Huber & Anders 2014) per treatment category and for biochar overall. Models used rarefied genus counts with all classified genera for normalization, ‘poscounts’ size factors, local dispersion fits, block and depth covariates (plus year for particle size), and default independent and Cook’s outlier filtering, with BH adjustments within treatment category and contrast type. Code and text were revised with the coding language model *opus v5*.*5* (Anthropic 2026) with supervision and validation, and authors take responsibility for content.

## Results

### Physical and chemical

Both compost and biochar separately lowered bulk density (D_b_), and biochar-compost raised SOM and P (**Fig. 1**). Compared to unamended soils, D_b_ was lower with compost by 0.209±0.099 g cm^-3^ (*t*_229.9_ =-2.10, *p*=0.037) and biochar alone by 0.302±0.099 g cm^-3^ (*t* _229.9_ =-3.04, *p*=0.0026), with weaker biochar effects at depth by 0.064±0.024 g cm^-3^ per cm (*t* _229.9_ =2.70, *p*=0.0074). Biochar-compost increased SOM by 5.5±1.9% (*t* _229.9_ =2.95, *p*=0.0036) and available P by 20.7±7.0 ppm (*t* _229.9_=2.95, *p*=0.0035) relative to unamended soils, whereas pH differed only marginally significantly among treatments (*F* _3,229.9_=2.13, *p*=0.097).

Among particle sizes, P was lower with >1 mm biochar than with compost alone by 15.9±5.7 ppm (*t* _686.9_=-2.80, *p*=0.0052), and Mg decreased with >1 mm biochar-compost by 37±14 ppm (*t* _686.9_=-2.66, *p*=0.0079) and with <1 mm biochar-compost by 34±14 ppm (*t* _686.9_ =-2.41, *p*=0.016). Deeper soils also had higher D_b_ by 0.026±0.011 g cm^-3^ per cm (*t* _686.9_=2.27, *p*=0.024) and lower SOM by 0.59±0.17% per cm (*t* _686.9_=-3.56, *p*<0.001).

**Fig 1.**
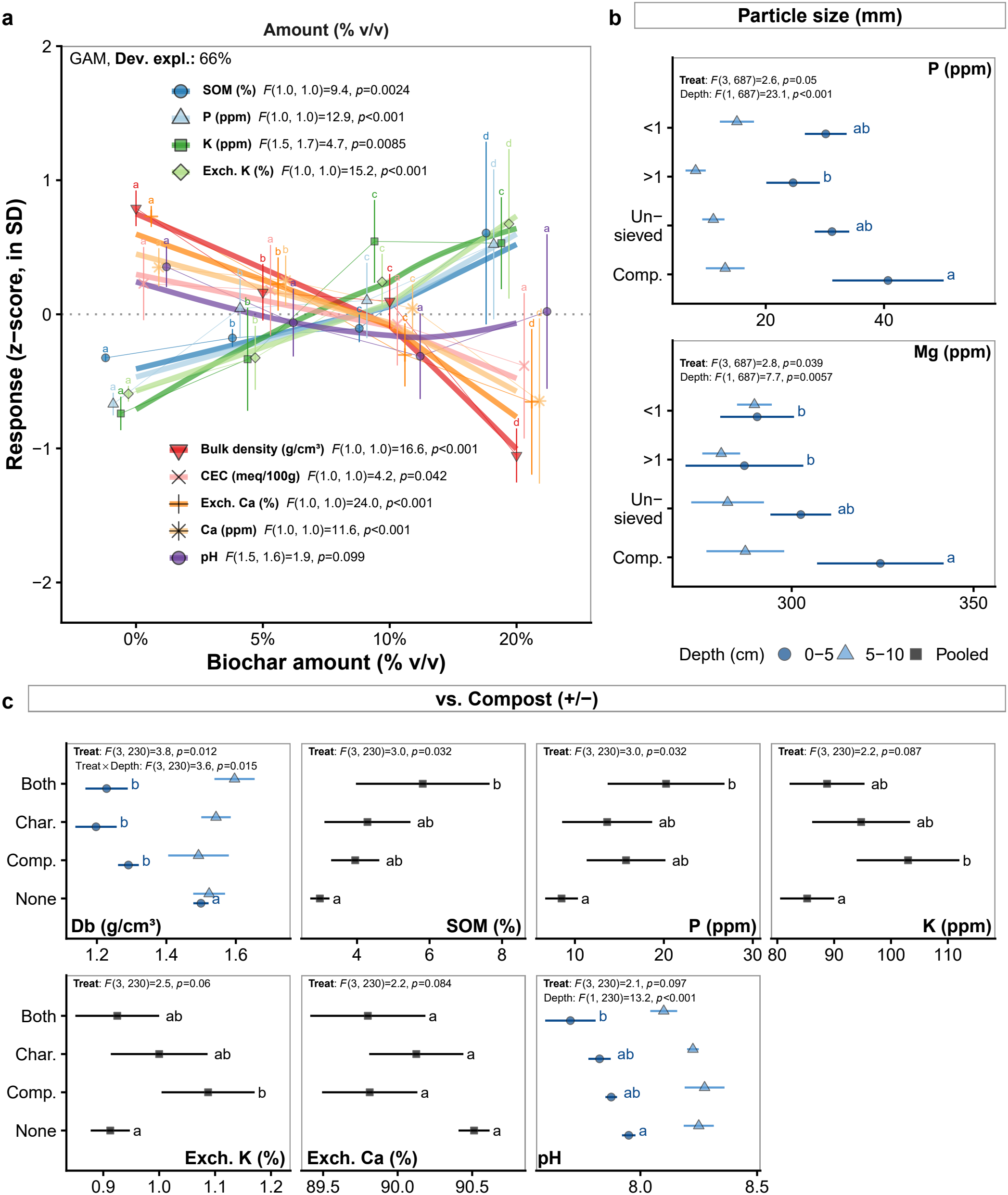
Biochar effects on urban soil nutrients. Significant biochar treatment effects on urban soil chemistry (MSU-DPFLI, Detroit, MI) using one GAM per treatment category and response variable scale. (**a**) Amount: *z*-scores (mean=0 dotted line, SD=1) across means±SE across samples (points; dark blue circles, SOM; light blue triangles, P; dark green squares, K; light green diamonds, exchangeable K; red inverted triangles, D_b_; pink crosses, CEC; orange plus signs, exchangeable Ca; light orange asterisks, Ca; purple circles, pH), smooth trends (thick lines), and pairwise contrast letters using *emmeans*. (**b**) Particle size and (**c**) *vs*. compost (None=unamended, Comp.=compost-only, Char.=biochar-only, Both=5% biochar-compost; Unsieved, >1, and <1=biochar-compost particle sizes, mm): means±SE by depth where significant (dark blue circles, 0-5 cm; light blue triangles, 5-10 cm; black squares, pooled), and pairwise treatment contrast letters (*p*<0.05).

Biochar application rate had the largest effects, lowering D_b_, Ca, and CEC but raising SOM, P, and K. D_b_ decreased with 5% by 0.089±0.022 g cm^-3^ (*t* _249.9_ =-4.07, *p*<0.001), 10% by 0.178±0.044 g cm^-3^ (*t* _249.9_ =-4.07, *p*<0.001), and most with 20% biochar co-application by 0.356±0.087 g cm^-3^ (*t* _249.9_ =-4.07, *p*<0.001). SOM increased with 20% biochar by 14.4±4.7% (*t* _249.9_=3.06, *p*=0.0024) and available P by 35.1±9.8 ppm (*t* _249.9_=3.60, *p*<0.001), K increased with both 10% by 51±18 ppm (*t* _249.9_=2.88, *p*=0.0043) and 20% biochar by 77±25 ppm (*t* _249.9_=3.11, *p*=0.0021), and exchangeable K increased with only 20% biochar by 1.65±0.42% (*t* _249.9_=3.90, *p*<0.001), with weaker effects at depth by 0.22±0.10% per cm (*t* _249.9_=-2.20, *p*=0.029). In contrast, 20% biochar lowered Ca by 941±276 ppm (*t* _249.9_=-3.41, *p*<0.001) and CEC by 1.77±0.87 meq 100 g^-1^ (*t* _249.9_ =-2.05, *p*=0.042), and exchangeable Ca decreased with 10% by 1.89±0.39% (*t* _249.9_=-4.90, *p*<0.001) and 20% biochar by 3.77±0.77% (*t* _249.9_=-4.90, *p*<0.001). Mg did not change with application rate, and pH increased more with depth at 20% biochar by 0.043±0.018 per cm (*t* _249.9_=2.32, *p*=0.021), *i*.*e*. surface pH was lower relative to subsurface pH.

### Biological

#### Composition

Biochar application rate and biochar *vs*. compost treatments changed bacterial composition, and particle size changed fungal dispersion. Bacterial composition shifted with biochar application rate (*R*^2^=0.13, *F*_3,19_=1.28, *p*<0.001), interactively with depth (*R*^2^=0.14, *F* _3,19_ =1.39, *p*=0.029), and among biochar *vs*. compost treatments (*R*^2^=0.12, *F* _3,20_ =1.12, *p*=0.036) without unequal dispersion, whereas fungal composition differed only marginally significantly among particle size treatments (*R*^2^=0.11, *F* _3,24_ =1.28, *p*=0.081), partly through unequal dispersion (**Fig. 2**). Multivariate dispersion (PERMDISP) differed among biochar amounts for bacteria (*F* _3,23_=2.59, *p*<0.001), and was unequal among particle size treatments for fungi (*F* _3,28_=3.36, *p*=0.03). Composition also varied with depth in all treatment categories. Bacterial composition across application rates aligned most with SOM (*r* ^2^=0.53, *p*=0.004), Ca (*r* ^2^=0.5, *p*=0.019), and CEC (*r* ^2^=0.42, *p*=0.036), and fungal composition under particle size with exchangeable K (*r* ^2^=0.46, *p*=0.003), SOM (*r* ^2^=0.45, *p*=0.001), and P (*r* ^2^=0.42, *p*=0.001) (all significant vectors in **Fig. 2**). Microbial alpha diversity (richness) and sample dispersion (distance to treatment × depth centroids; GAMs) remained consistent across treatments.

**Fig 2.**
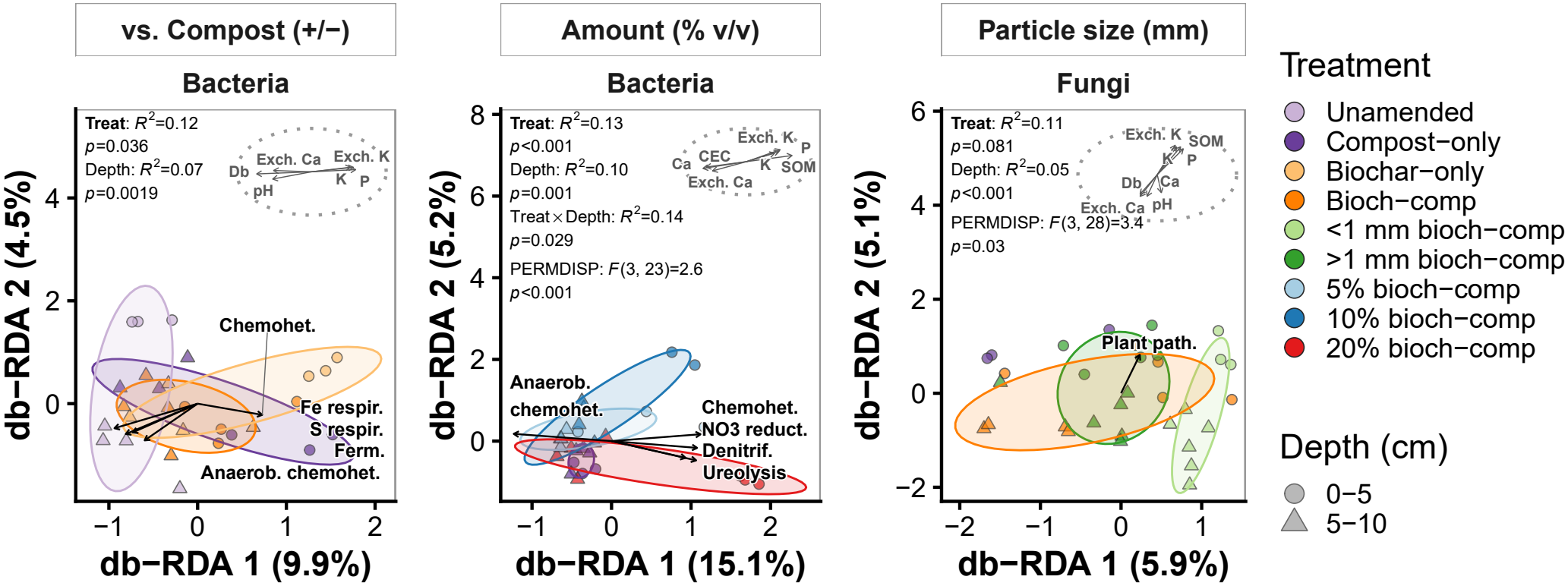
Biochar effects on urban soil microbial composition. Significant biochar treatment effects on an urban soil (MSU-DPFLI, Detroit, MI; db-RDA of Bray-Curtis distances; axes, % variation): 68% treatment ellipses (colors: purple, compost-only; orange, biochar-compost; light and dark green, <1 and >1 mm; light and dark blue, 5 and 10%; red, 20%; circles, 0-5 cm; triangles, 5-10 cm), envfit arrows (*p*<0.05; length ∝ *r* ) for the five strongest functional groups (black) and soil variables (grey inset; dotted circle, *r* =1; D_b_, bulk density), and PERMANOVA and PERMDISP tests (corner).

#### Assembly

Biochar and compost treatments significantly affected the relative importance of community assembly processes, with drift, dispersal limitation, and homogeneous selection dominating. Bacterial community assembly was dominated by ecological drift (68% of pairwise turnover), followed by dispersal limitation (20%) and homogeneous selection (11%), with little heterogeneous selection or homogenizing dispersal (1.7% combined; **Fig. 3** a). At 0-5 cm, dispersal limitation increased relative to each treatment category’s reference, with unsieved biochar-compost increasing by 10.9% (*F* _3,226_=4.1, *p*=0.0073), biochar alone by 18.9% (*F* _3,101_=20.1, *p*<0.001), and 20% biochar by 16.2% (*F* _1.0,1.0_=13.7, *p*<0.001). Biochar alone also decreased drift by 24% (*F* _3,101_=12.7, *p*<0.001) and homogeneous selection by 3.3% (*F* _3,101_=5.0, *p*=0.003), and increased homogenizing dispersal by 7.4% (*F* _3,101_=16.2, *p*<0.001). Among particle sizes, >1 mm biochar-compost increased homogenizing dispersal by 0.9% (*F* _3,226_=4.0, *p*=0.0085), and <1 mm biochar-compost increased heterogeneous selection by 2.4% (*F* _3,226_=8.9, *p*<0.001). Fungal assembly was also dominated by drift (50%), but with more dispersal limitation (35%) and heterogeneous selection (4%) than bacterial assembly (**Fig. 3** b). Biochar shifted fungal assembly from drift toward selection and dispersal limitation at 0-5 cm. Among particle sizes, <1 mm biochar-compost decreased drift by 42.7% (*F* _3,100_=18.6, *p*<0.001) and increased homogeneous selection by 22.9% (*F* _3,100_=6.0, *p*<0.001), while >1 mm biochar-compost increased dispersal limitation by 36.3% (*F* _3,100_=15.8, *p*<0.001). 20% biochar decreased drift by 14% (*F* _1.9,2.0_=4.9, *p*=0.014) and increased homogeneous selection by 16.7% (*F* _1.9,2.0_=20.0, *p*<0.001) and heterogeneous selection by 4% (*F* _1.0,1.0_=6.3, *p*=0.013), and biochar-compost raised heterogeneous selection *vs*. unamended soils by 15.2% (*F* _3,67_=15.1, *p*<0.001).

**Fig 3.**
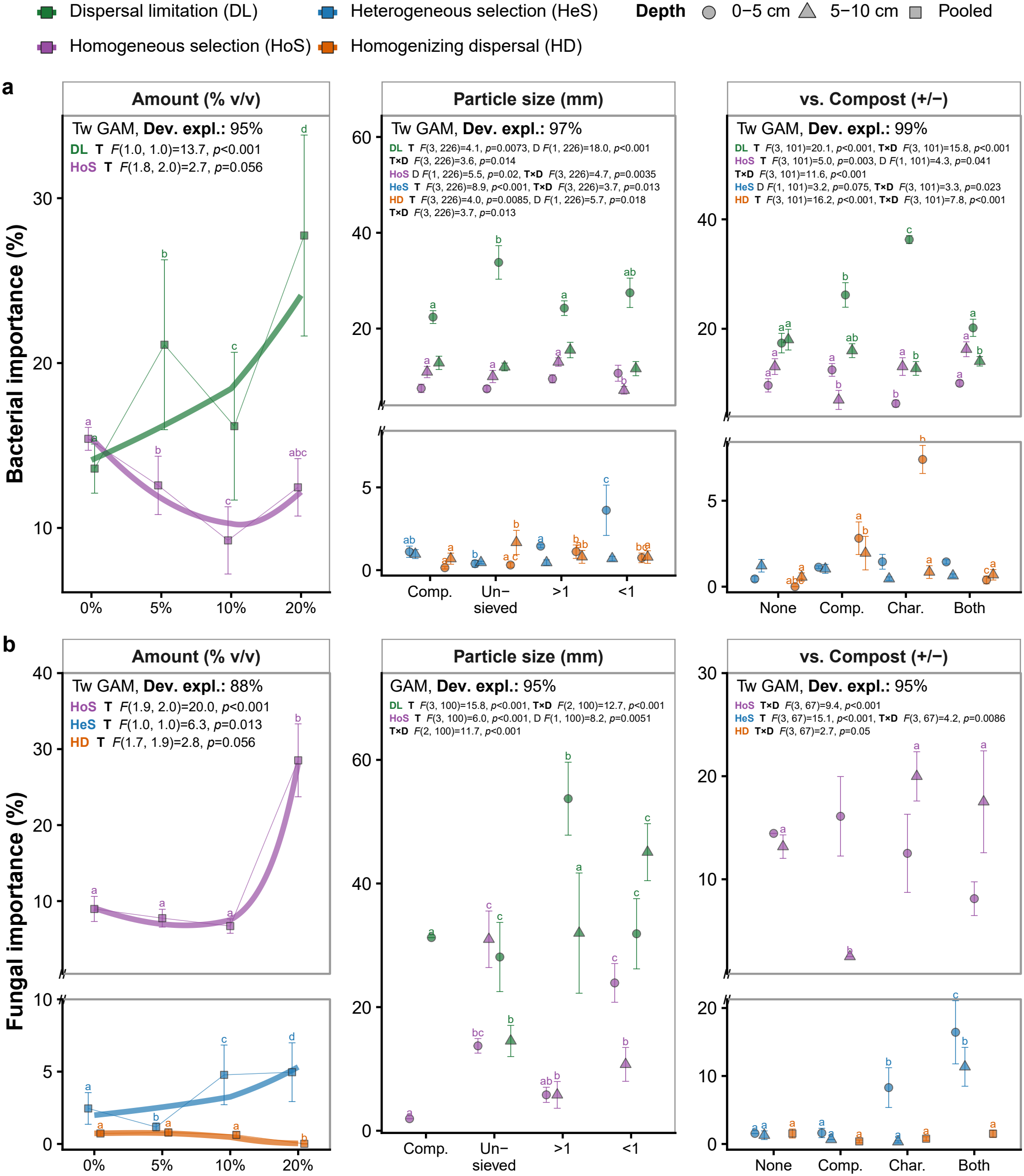
Bacterial and fungal community assembly processes in biochar-amended urban soils. Processes with significant treatment effects on (**a**) bacteria and (**b**) fungi (MSU-DPFLI, Detroit, MI; iCAMP; one GAM per domain and treatment category; drift not shown): means±SE across samples (points; dark green, dispersal limitation; purple, homogeneous selection; blue, heterogeneous selection; orange, homogenizing dispersal; circles, 0-5 cm; triangles, 5-10 cm; squares, pooled where depth terms were not significant; letters, pairwise differences) and Amount smooths at mean depth (lines); axes break (//) above processes <5%; tests in each panel (T, treatment; D, depth; T×D, interaction).

#### Functional groups

Biochar co-additions, especially at 20%, increased predicted heterotrophic and N-cycling bacterial functions along with fungal soil saprotrophs. Of 14 common bacterial functions predicted from taxonomy, 20% biochar increased predicted chemoheterotrophy by 17.6±5.0% (*t* _505.2_=3.49, *p*<0.001), nitrate reduction by 2.77±0.60% (*t* _505.2_=4.59, *p*<0.001), denitrification by 1.44±0.50% (*t* _505.2_=2.88, *p*=0.0042), ureolysis by 1.94±0.52% (*t* _505.2_=3.73, *p*<0.001), methylotrophy by 1.01±0.34% (*t* _505.2_=3.00, *p*=0.0028), and predatory or exoparasitic bacteria by 0.74±0.33% (*t* _505.2_=2.23, *p*=0.026), but decreased anaerobic chemoheterotrophy by 6.0±2.5% (*t* _505.2_=-2.34, *p*=0.02), mostly at 0-5 cm (**Fig. 4 a**). Relative to unamended soils, compost alone increased 0-5 cm fermentation n.s. and methylotrophy by 0.79±0.31% (*t* _379_=2.56, *p*=0.011), and biochar alone increased predatory or exoparasitic bacteria by 0.98±0.33% (*t* _379_=2.94, *p*=0.0035), while unsieved biochar-compost together increased dark hydrogen oxidation under particle size treatments by 1.50±0.40% (*t* _865.9_=3.80, *p*<0.001). Among 11 common fungal lifestyles, pollen saprotrophs increased with large biochar particles by 1.27±0.40% (*t* _865.9_=3.14, *p*=0.0018), whereas mycoparasites decreased with >1 mm biochar-compost by 14.5±5.8% (*t* _865.9_=-2.50, *p*=0.013), with <1 mm biochar-compost by 15.6±6.7% (*t* _865.9_=-2.34, *p*=0.019), and with unsieved biochar-compost by 15.2±5.9% (*t* _865.9_=-2.59, *p*=0.0099) under particle size treatments. Soil fungal saprotrophs increased with 20% biochar by 31±11% (*t* _505.2_=2.85, *p*=0.0045), and animal parasites decreased with compost alone by 17.5±5.4% (*t* _379_=-3.27, *p*=0.0012), with biochar alone by 17.4±5.4% (*t* _379_=-3.25, *p*=0.0012), and with biochar-compost by 17.7±5.6% (*t* _379_=-3.13, *p*=0.0019) (**Fig. 4** c).

**Fig 4.**
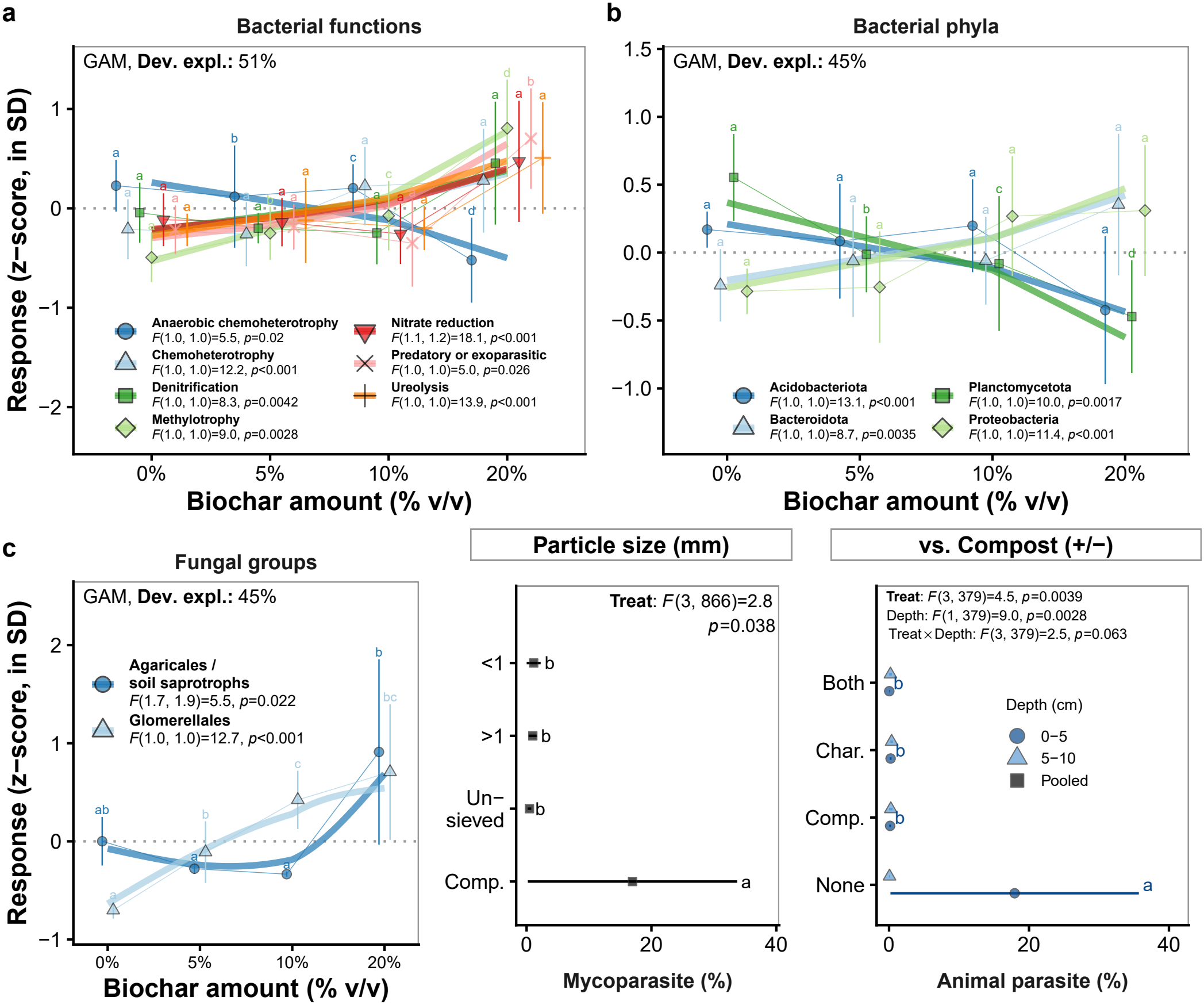
Microbial functions and taxon responses to urban biochar. Significant treatment effects (MSU-DPFLI, Detroit, MI; one GAM per treatment category and response group) on (**a**) predicted bacterial functions, (**b**) dominant bacterial phyla, and (**c**) fungal orders (among the eight most abundant) with biochar application rate, and fungal parasites. Amount panels show *z*-scores (mean=0 dotted line, SD=1; points, means±SE across samples; letters show pairwise differences). In (**a**), dark blue circles show anaerobic chemoheterotrophy, light blue triangles chemoheterotrophy, dark green squares denitrification, light green diamonds methylotrophy, red inverted triangles nitrate reduction, pink crosses predatory or exoparasitic bacteria, and orange plus signs ureolysis. In (**b**), dark blue circles show Acidobacteriota, light blue triangles Bacteroidota, dark green squares Planctomycetota, and light green diamonds Proteobacteria. In (**c**), dark blue circles show Agaricales (similar pattern for soil saprotrophs) and light blue triangles Glomerellales. Other panels show means±SE (dark blue circles, 0-5 cm; light blue triangles, 5-10 cm; black squares, pooled) with pairwise contrast letters (*p*<0.05).

Biochar shifted dominant phyla mainly at 20%, increasing Proteobacteria by 13.4±4.0% (*t* _300.9_=3.38, *p*<0.001) and Bacteroidota by 14.9±5.0% (*t* _300.9_=2.95, *p*=0.0035) but decreasing Acidobacteriota by 15.2±4.2% (*t* _300.9_=-3.62, *p*<0.001) and Planctomycetota by 4.4±1.4% (*t* _300.9_=-3.16, *p*=0.0017), mostly at 0-5 cm (**Fig. 4** b). Among the eight most abundant fungal orders, mushroom-forming Agaricales increased with 20% biochar by 28±11% (*t* _300.9_=2.50, *p*=0.013) and Glomerellales increased with application rate, most at 20% by 2.75±0.77% (*t* _300.9_=3.56, *p*<0.001) (**Fig. 4** c). Hypocreales decreased with compost by 17.4±6.1% (*t* _208.9_=-2.88, *p*=0.0044), biochar by 13.7±6.1% (*t* _208.9_=-2.26, *p*=0.025), and biochar-compost by 17.6±6.3% (*t* _208.9_=-2.77, *p*=0.0061) relative to unamended soils, and Helotiales increased with depth under compost alone by 1.67±0.64% per cm (*t* _208.9_=2.61, *p*=0.0097).

**Table 1.** Responsive modules. Top 10 named genera (fungi †; % of module abundance) of responsive modules (MSU-DPFLI, Detroit, MI) and surface responses *vs*. each treatment category’s reference (↑ higher, ↓ lower; *p*<0.05).

| **Network module** | **Response** | **Top genera (% of module)** |
|---|---|---|
| **Copiotroph** | ↑ <1<br>(Particle size) | *Dominikia*† (26%), *Flavobacterium* (11%), *Coprinopsis*† (7%), *Paraphoma*† (7%), *Chthoniobacter* (7%), *Pseudomonas* (6%), *Sphingomonas* (4%), *Fusarium*† (3%), *Ohtaekwangia* (3%), *Ferruginibacter* (3%) |
| **Oligotroph** | ↑ <1<br>(Particle size)<br>↓ 5%, 10%, 20%<br>(Amount) | *Flavisolibacter* (28%), *Bryobacter* (20%), *Cercospora*† (15%), *Sumerlaea* (9%) |

#### Networks

The signed co-occurrence network (96 bacterial and 33 fungal genera, 835 edges, 283 negative) was more modular (*Q*=0.42 vs. 0.31 random, *p*=0.001) and clustered (*C*=0.48 vs. 0.07 random, *p*=0.001) than random networks, and negative edges linked different modules more often than expected (100% *vs*. 83% with shuffled modules, *p*=0.001); two modules responded to treatments and two only marginally significantly (**Fig. 5**, Table 1). A copiotroph module, including *Pseudomonas, Flavobacterium*, and *Variovorax* with fungal *Plectosphaerella*, increased with small biochar particles at 0-5 cm by 37±16% (*t* _109.4_=2.32, *p*=0.022). An oligotroph module of acidobacterial *Bryobacter* and Sub-group 10 decreased with biochar amount, most with 20% biochar by 6.2±2.2% (*t* _112.2_=-2.87, *p*=0.0049), whereas a nitrifier module grouping nitrifying *Nitrospira* and iron-reducing *Geobacter* with arbuscular mycorrhizal *Glomus* (*F* _1,112.2_=3.42, *p*=0.067) and an N-fixer module (*F* _1.8,112.2_=3.06, *p*=0.078) varied only marginally significantly with biochar amount.

The whole network’s degree distribution was consistent with a power law (bootstrap *p*=0.35, *α*=5.65; Clauset, Shalizi & Newman 2009) but fit no better than exponential (Vuong *R*=-1.64, *p*=0.1) or lognormal distributions (Vuong *R*=-0.98, *p*=0.33), as in most empirical networks (Broido & Clauset 2019). Likewise, 89% of sample sub-networks were consistent with power laws (bootstrap *p*≥0.05), but power laws fit worse than exponential and lognormal distributions in 21% and 7% of sub-networks, and better in none (Vuong *p*<0.05).

Biochar treatments tended to make microbial networks sparser. With increasing biochar amount, sub-networks lost connections and hubs: at 20% biochar, mean degree decreased by 3.2±1.2 (*t* _235.4_=-2.74, *p*=0.0065), clustering by 0.157±0.066 (*t* _235.4_=-2.39, *p*=0.018), centralization by 0.115±0.037 (*t* _235.4_=-3.15, *p*=0.0019), and robustness by 0.102±0.035 (*t* _235.4_=-2.89, *p*=0.0042), whereas mean path length increased by 0.187±0.066 (*t* _235.4_=2.85, *p*=0.0048). Biochar alone had similar effects relative to unamended soil, lengthening mean path length by 0.295±0.095 (*t* _115.5_=3.11, *p*=0.0023) and raising modularity by 0.167±0.066 (*t* _115.5_=2.52, *p*=0.013), while lowering clustering by 0.188±0.082 (*t* _115.5_=-2.28, *p*=0.024) and robustness by 0.120±0.053 (*t* _115.5_=-2.29, *p*=0.024). Centralization decreased with both compost alone by 0.138±0.053 (*t* _115.5_=-2.62, *p*=0.0098) and biochar alone by 0.150±0.051 (*t* _115.5_=-2.94, *p*=0.004), but not with biochar-compost together.

Differential abundance tests identified only fungal genera, mostly copiotroph-module members and saprotrophs. Under particle size treatments, copiotroph-module saprotroph *Coprinopsis* was higher with <1 mm by 11.1±3.1 log_2_-fold (*q*=0.015) and unsieved biochar-compost by 9.3±3.1 log_2_-fold (*q*=0.029) than with compost alone, whereas sapro-trophic *Linnemannia* was lower with <1 mm biochar-compost by 6.0±1.8 log_2_-fold (*q*=0.018). Relative to unamended soils, copiotroph-module pathogen *Fusarium* was lower with compost alone by 7.8±2.3 log_2_-fold (*q*=0.0061), biochar alone by 5.8±2.2 log_2_-fold (*q*=0.034), and biochar-compost by 9.3±2.3 log_2_-fold (*q*<0.001), whereas *Coprinopsis* was higher with biochar alone by 7.3±2.8 log_2_-fold (*q*=0.034). Saprotrophs outside responsive modules shifted both ways: soil saprotrophs *Mortierella* decreased with biochar overall by 2.81±0.91 log_2_-fold (*q*=0.025), most so with biochar-compost by 4.5±1.4 log_2_-fold (*q*=0.0063), and potential saprotroph *Linnemannia* decreased with biochar alone by 3.8±1.5 log_2_-fold (*q*=0.034), whereas dung saprotroph *Preussia* increased with compost alone by 18.4±2.4 log_2_-fold (*q*<0.001), biochar alone by 20.0±2.3 log_2_-fold (*q*<0.001), and biochar-compost by 17.3±2.4 log_2_-fold (*q*<0.001), and arbuscular mycorrhizal *Rhizophagus* increased with compost alone by 7.3±2.4 log_2_-fold (*q*=0.012).

**Fig 5.**
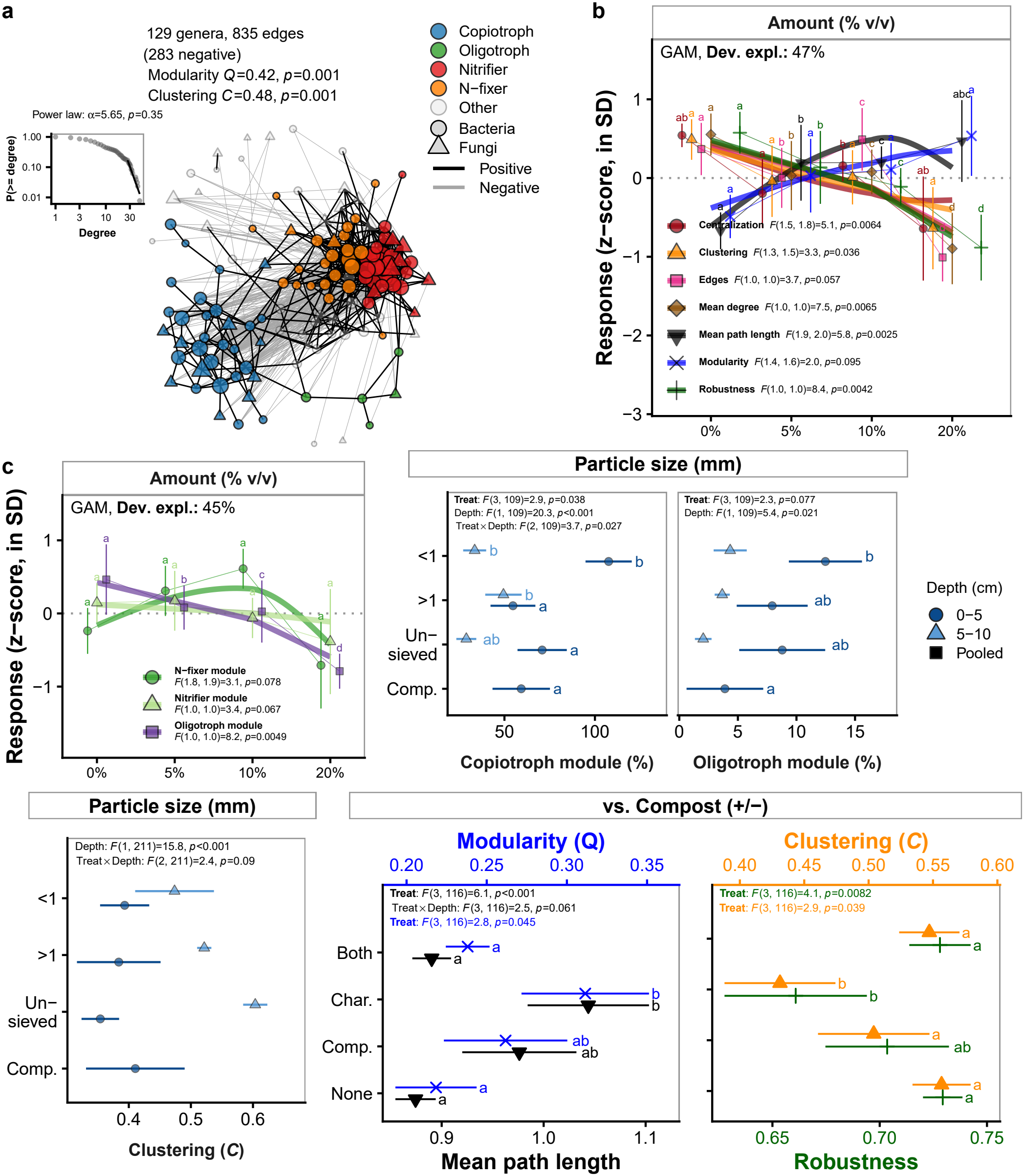
Co-occurrence network. (**a**) Signed bacterial (circles) and fungal (triangles) genus co-occurrence network (MSU-DPFLI, Detroit, MI; CLR correlations; black, positive; grey, negative edges); modules colored (blue, copiotroph; green, oligotroph; red, nitrifier, and orange, N-fixer, marginally significant; **Table 1**; faded grey, other), nodes sized by degree; modularity (*Q*) and clustering (*C*) of positive edges *vs*. random networks (corner); inset, cumulative degree distribution with power-law fit (log-log). (**b**) Sub-network properties and (**c**) module abundances with treatment effects; Amount as *z*-scores (as in **Fig. 4**), other treatment categories as means±SE (as in **Fig. 1**), with similar *vs*. compost properties paired (second on the colored top axis).

Neither invertebrate richness nor dispersion differed among particle size treatments, but showed possible trends in differing composition. Invertebrate composition was mainly similar among treatments (*R*^2^=0.04, *F* =0.82, *p*=0.86), along with similar multivariate dispersion (*F* _3,60_=0.93, *p*=0.44). However, compositional trends appeared to align with the proportions of Araneae (*r* ^2^=0.34, *p*=0.001), Coleoptera (*r* ^2^=0.18, *p*=0.011), and Acari (*r* ^2^=0.09, *p*=0.038) (**Fig. 6**), but remained independent of soil variables (envfit *p*>0.05). Proportions of the main orders (Collembola, Hymenoptera, Acari, Araneae, Coleoptera, and Hemiptera) were consistent among treatments.

**Fig 6.**
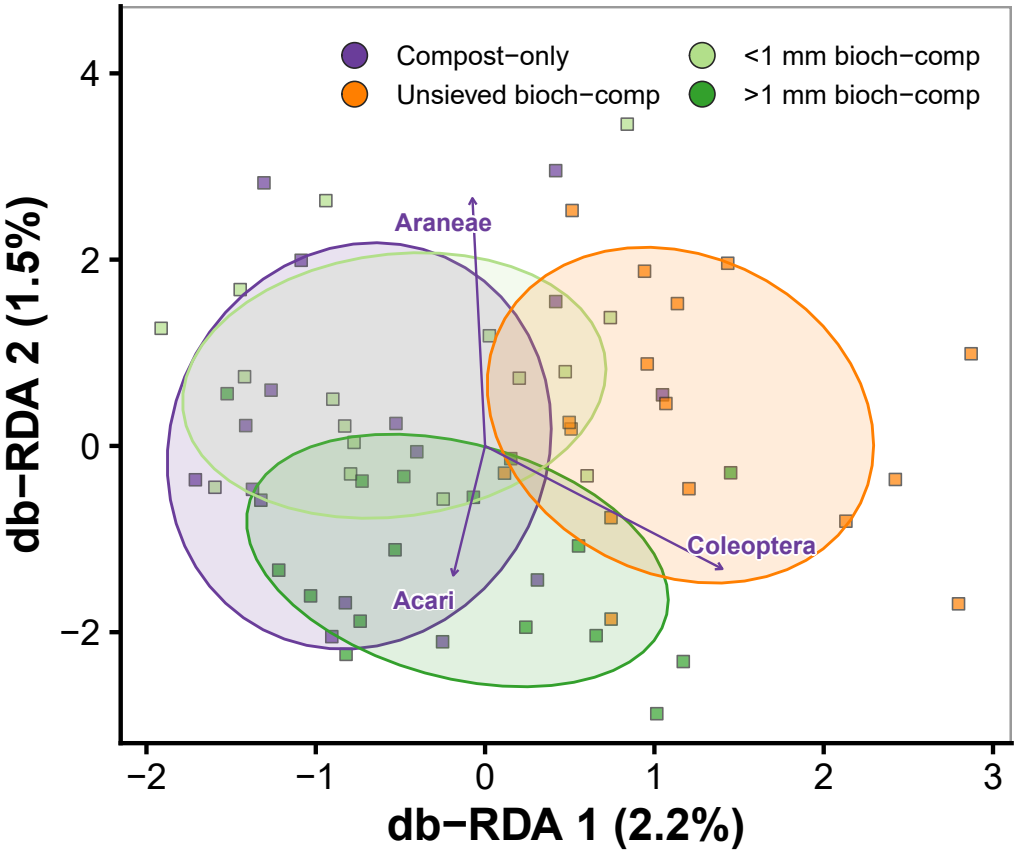
Invertebrate composition. Particle size treatments (MSU-DPFLI, Detroit, MI; squares, traps colored by treatment: purple, compost-only; orange, biochar-compost; light and dark green, <1 and >1 mm; db-RDA as in **Fig. 2**; treatment n.s.): 68% ellipses and envfit arrows for orders (*p*<0.05).

## Discussion

Taken together, these results show that various short-term biochar amendments improved post-construction urban soils, overall lowering bulk density and increasing SOM, P, and K, while reorganizing rather than enriching soil microbial communities. Consistent with the first hypothesis, biochar-compost differed from either amendment alone: biochar alone lowered bulk density more than compost alone, only biochar-compost raised P and suppressed bacterial dispersal limitation, and bacterial composition differed among treatments. Partly consistent with the second hypothesis, particle size did not change soil properties but did change biota, where smaller <1 mm particles increased topsoil copiotrophs and decomposers, fungal assembly shifted from drift toward more homogeneous selection and dispersal limitation, and fungal communities were more heterogeneous with biochar of either size than with compost alone. Results were consistent with the third hypothesis, where higher biochar amounts lowered bulk density and raised SOM, but also lowered Ca and CEC, made bacteria more heterogeneous (dispersed), but biochar effects were not mediated by changes in species richness. Biologically, biochar overall increased fast-growing copiotrophs and decreased oligotrophs, increased bacterial dispersal limitation and fungal homogeneous selection, and increased potential N-cycling and decomposer functions.

Compared with compost, biochar helped reduce bulk density, but additional compost helped to raise SOM, P, lower pH, and balance microbial assembly processes and biodiversity. Biochar alone lowered bulk density more than compost alone, reflecting lower biochar density (Werdin *et al*. 2020) and irregular particle shapes (Brewer *et al*. 2014), and also mirrored the reduced penetration resistance under roto-tilling at the same site (Edwards, Medina, and Asker 2023), potentially suggesting an urban soil compaction solution with less disturbance but a slightly higher cost (∼2.7 USD m^-2^ [0.25 USD ft^-2^] more, *pers. comms*.). Together with compost, P and SOM increased, suggesting biochar retains compost P without supplying it (Zhang *et al*. 2015; Small *et al*. 2019). Microbially, biochar alone shifted topsoil bacterial assembly away from homogeneous selection toward dispersal limitation, and lowered network clustering and robustness, which may be due to more fragmented microhabitats among biochar particles (Quilliam *et al*. 2013), and was restored by mixing in compost. These shifts ultimately lowered potentially pathogenic *Fusarium* and saprotrophic *Mortierella* while favoring decomposer *Coprinopsis*, pointing to a need for future work elaborating on potential disease pressure, protection, and/or decomposition functions, especially as P-scavenging Glomerales, including arbuscular mycorrhizal fungi (Mowrer *et al*. 2022), did not respond to higher P.

Interestingly, biochar particle size changed biota more than soil properties. Contrary to initial hypotheses, biochar particle size did not affect bulk density, likely because 0.1-2 mm remains coarse compared with compacted clays (Liao & Thomas 2019; Zhang, Amonette & Flury 2021). Notably, small particles in topsoils did favor decomposer fungus *Coprinopsis* and a copiotroph module of rhizosphere-associated bacteria *Pseudomonas, Flavobacterium*, and *Variovorax*, likely associated with higher surface area and contact with compost and roots with potentially more accessible pools of labile C and nutrients. Fungi were generally more sensitive to particle size changes, being more heterogeneous with either particle size, shifting assembly from drift toward homogeneous selection with smaller particle sizes, and being more dispersal limited among larger particle sizes. Taken together, alongside larger particles diluting P and Mg, these findings suggest that biochar filters fungi more deterministically than bacteria, which were only dispersal limited among all unsieved particles in topsoil. Invertebrates were relatively consistent among particle size treatments, although trends toward mite groups enriched among larger biochar particles and spider groups enriched among smaller particles may suggest indirect trophic effects on each’s prey populations. Overall, invertebrate results aligned with relatively inconsistent faunal responses to biochar (Domene 2016), lags behind soil and microbial changes (Hylander & Ehrlén 2013), or negative effects of dissolved biochar derivatives (Gomez-Eyles *et al*. 2011; Godlewska, Ok, and Oleszczuk 2021), although invertebrate biochar studies otherwise remain notably sparse (Singh *et al*. 2022; Zhu *et al*. 2023).

Biochar application rate had the largest effects among treatment categories, increasingly improving soil chemistry but with a trade-off in diluted Ca and CEC. Bulk density decreased with application rate, as in other studies (Blanco-Canqui 2017; Wong, Levy-Sakin, and Kwok 2018). Biochar also increased SOM, P, and K (Saarnio *et al*. 2018; Siedt *et al*. 2021), which may be concentrated from the compost co-additions (*ca*. 2% N and 10:1 C:N; Ryals *et al*. 2016). Consistent with higher nutrient concentrations, application rate also increased bacterial heterogeneity and favored copiotrophs, such as those in Bacteroidota (Shiu *et al*. 2022), decreased oligotrophic acidobacterial *Bryobacter*, and increased chemoheterotrophy and N-cycling functions, overall suggesting faster nutrient cycling and/or SOM turnover. This may also be consistent with bacterial assembly shifting toward dispersal limitation and less network clustering at 20% application, as copiotrophs may be able to grow more easily on available nutrients in a micro-habitat. In contrast, fungal assembly shifting from drift toward homogeneous selection at 20% application may again support relatively more deterministic assembly at these spatial scales compared to bacteria, such as of Agaricales and saprotrophic decomposers (Tanesaka, Masuda, and Kinugawa 1993; Floudas *et al*. 2020). Across treatments, network analyses grouped nitrifying and iron-reducing bacteria with arbuscular mycorrhizal *Glomus*, suggesting potentially cooperative cross-domain functional guilds, supported by emerging hyphosphere research (Ravnskov, Nybroe, and Jakobsen 1999; Jiang *et al*. 2021; Zhang *et al*. 2022; Nguyen 2023; Wang *et al*. 2023; Vieira *et al*. 2025; Vaishnav & Jansa 2026).

Overall, this study shows that biochar co-amendments can offer physical and chemical benefits for urban agricultural soils, likely greatest in low-nutrient soils (Crane-Droesch *et al*. 2013; Noyce *et al*. 2015; Jeffery *et al*. 2017; Deng *et al*. 2026), reorganizing microbial community functions and underlying assembly processes. This study newly emphasizes how various biochar usage strategies affect not simply chemical but also biological and fundamental ecological processes from soil physical particles to microorganisms and invertebrates, supporting future work on mechanisms behind small-scale biochar adoption. Holistic analyses of biochar adoption within urban social-ecological systems will offer more realistic perspectives on biochar’s wider role in managing urban lands under turnover.

## Acknowledgements

Thanks to the Vandermeer and Perfecto labs and Blesh labs for advice; MSU-DPFLI field site partners and interns for helpful discussions and field assistance; Keep Growing Detroit for curating seeds, public engagement, and urban agriculture knowledge; Cadillac Urban Gardens for initial discussion; and Sweetland peer review group for early draft feedback.

